# Utilizing citizen science to model the distribution of *Aedes aegypti* ([Diptera]: [Culicidae]) in West Africa

**DOI:** 10.1101/2022.01.14.476313

**Authors:** Elizabeth A. Freeman, Elizabeth J. Carlton, Sara Paull, Samuel Dadzie, Andrea Buchwald

**Author notes:** **Corresponding author:** Elizabeth Freeman.

## Abstract

In a rapidly urbanizing region such as West Africa, *Aedes* mosquitoes pose an emerging threat of infectious disease that is compounded by limited vector surveillance. Citizen science has been proposed as a way to fill surveillance gaps by training local residents to collect and share information on disease vectors. Increasing citizen science efforts can begin to bridge the gaps in our current knowledge of *Aedes* distribution while engaging locals with mosquito control and public health efforts. Understanding the distribution of disease vectors in West Africa can inform researchers and public health officials on where to conduct disease surveillance and focus public health interventions. We aimed to compare citizen science data to published literature observations of *Aedes* mosquitoes and to quantify how incorporating citizen science changes our understanding of *Aedes* mosquito distribution in West Africa. We utilized citizen science data collected through NASA’s GLOBE Observer mobile phone application and data from a previously published literature review on *Aedes* mosquito distribution to examine the contribution of citizen science to understanding the distribution of *Ae. aegypti* in West Africa using Maximum Entropy modeling. Combining citizen science and literature-derived observations improved the fit of the model compared to models created by each data source alone, but did not alleviate location bias within the models, likely due to lack of widespread observations. Understanding *Ae. aegypti* distribution will require greater investment in *Aedes* mosquito surveillance in the region, and citizen science should be utilized as a tool in this mission to increase the reach of surveillance.

## Introduction

Arboviral diseases transmitted by *Aedes* mosquitoes are an emerging threat to the health of populations globally (Leta et al. 2018, Ryan et al. 2019), however there are data gaps regarding the distribution of key mosquito vectors in many regions, such as West Africa. West Africa has experienced multiple outbreaks of Zika, dengue and chikungunya over the past 15 years, yet the distribution of *Aedes* vectors is poorly defined (Buchwald et al. 2020). Vector surveillance can help public health officials to better predict where and when *Aedes*-borne diseases will emerge, as disease follows patterns of vector abundance (Carlson et al. 2016, Messina et al. 2016, Leta et al. 2018). Global environmental models of *Aedes aegypti* and *Ae. albopictus* have been created (Kraemer et al. 2015, Kamal et al. 2019, Ryan et al. 2019), but these lack the fine-scale spatial resolution desired by decision-makers and are subject to climate bias from global collinearity of environmental variables (Dormann et al. 2013). In many regions around the world where systematic arbovirus surveillance infrastructure is lacking, there is a need for enhanced local surveillance in order to improve epidemic preparedness and early warning systems. Given recurrent yellow fever outbreaks and the increasing public health burden due to dengue and chikungunya, West Africa is a priority region for strengthening the public health entomology capacities around *Aedes* surveillance and control. Improved *Aedes* surveillance and modelling will provide essential information for the formulation of evidence-based vector control strategies and also improve preventive actions to protect human populations from the diseases transmitted by *Aedes* mosquitoes.

Citizen science has been proposed as a way to bolster local *Aedes* mosquito surveillance (Walther and Kampen 2017, de Souza Silva et al. 2018, Eritja et al. 2019, Amos et al. 2020). Citizen science is generally defined as members of a local or global community-many without formal training-participating in scientific activities, such as data collection and species surveillance (Kullenberg and Kasperowski 2016). Citizen science approaches have been used to enhance surveillance for multiple mosquito-borne diseases (Lwin et al. 2014, Hardy and Barrington 2017, Palmer et al. 2017, Richman et al. 2018). Utilizing citizen scientists can improve *Aedes* surveillance: residents may have preexisting knowledge of where mosquitoes reside in their community and can help scientists decide where to focus research (Mwangungulu et al. 2016). Participating in citizen science also benefits the community by increasing knowledge of mosquito-borne diseases and can spark behavioral changes to mitigate risk, such as engaging in household mosquito breeding site reduction (Toledo Romani et al. 2007, Stefopoulou et al. 2018, Asingizwe et al. 2020). However, the utility of citizen science programs can be doubtful due to challenges including recruitment and training of citizen scientists, as well as issues of data integrity (Johnson et al. 2018).

This study examines the value of incorporating data from NASA’s Global Learning and Observations to Benefit the Environment (GLOBE) Observer Mosquito Habitat Mapper (MHM) application (https://observer.globe.gov/toolkit/mosquito-habitat-mapper-toolkit) (Amos et al. 2020) to estimate the distribution of *Aedes* mosquitoes in West Africa. The GLOBE Program has regional offices around the world and conducts outreach to students and teachers; providing them curriculum and protocols for conducting citizen science-however people who do not receive training can also use the app. In 2018, the GLOBE Zika Education and Prevention Project led an effort in 30 countries to bolster MHM observations (Aïkpon 2019). The MHM was designed to address two goals; to categorize and map water sources used as mosquito breeding grounds and to understand the distribution of mosquitoes. The app primarily captures data on the larval stage of mosquito development, but additionally has users input information about other life stages present at the habitat (Amos et al. 2020). GLOBE Observer walks the user through a series of questions to identify habitat types and the genus of mosquitoes found. This process requires the user to have access to mosquito larvae sampling equipment and magnifying lens for their phone camera (Muñoz et al. 2020). From 2016 to 2019, there were 320,000 observations submitted in the app by 38,000 users (Amos et al. 2020).

In this study, we aim to evaluate the utility of the citizen science data for enhancing the characterization of environmental suitability for *Aedes* mosquitoes across West Africa. To do this, we examine citizen science observations from GLOBE MHM and data from a literature review to estimate distribution of *Ae. aegypti* in West Africa. We use Maximum Entropy modeling to evaluate the extent to which citizen science data can improve estimates of the spatial distribution of the primary *Aedes-borne* disease vector.

## Methods

We modeled distribution of *Ae. aegypti* mosquitoes in West Africa using two different data sources: i) observations extracted from GLOBE MHM, and ii) a compilation of the published literature containing scientific reports on *Aedes* surveillance from Buchwald et al. 2020 (Buchwald et al. 2020). There are multiple methods to model species distribution (Vezzani et al. 2005, Kearney et al. 2009, Richman et al. 2018), we chose Maximum Entropy modeling (Maxent) because it is well suited to the small numbers of observations, and presence-only data characterizing our dataset. Maxent modeling combines species observation data with key climate predictors of species presence to predict the probability of species presence (Phillips and DudÍk 2008). We compared the ranges for Ae. *aegypti* predicted by each data source alone and in combination to evaluate whether inclusion of citizen science data can improve estimates of species distribution in a region lacking data.

### Study Area

We defined the study area of West Africa as between the latitude of 20 and −30 and between the longitude of 20 and −10. This area encompasses Burkina Faso, Benin, Cameroon, Cape Verde, Chad, Côte d’Ivoire, Equatorial Guinea, Gabon, Gambia, Ghana, Guinea, Guinea Bissau, Liberia, Mali, Mauritania, Niger, Nigeria, Sao Tome and Principe, Sierra Leone, and Togo. These boundaries were chosen because they contain all *Aedes* presence points in both GLOBE MHM and the literature data used in the study.

### Mosquito Presence data

Data from GLOBE MHM were obtained via the GLOBE Observer’s Advanced Data Access Tool (https://datasearch.globe.gov/). We used data from the Mosquito Habitat Mapper and Mosquito Larvae protocols and searched by site name for all countries in our study area. Analysis was done in Rstudio (RCoreTeam 2021). Data were accessed on XXXX December 8^th^, 2020 and observations ranged from July 5th, 2017 to November 22nd, 2020. We selected observations that marked “true” for mosquito eggs, pupae, or adults or a non-zero larvae count for inclusion in further analysis. Most observations in the GLOBE dataset lacked genus and species data. Observations were filtered to include only unique locations through the removal of duplicates.

Mosquito observations from the published literature were collected as part of a previous study by Buchwald et al. covering the time period from 1999 through October 2018 (Buchwald et al. 2020). For this study, we restricted our main analysis to *Aedes* observations published from 2009-2018 to overlap with the time period of GLOBE MHM observations. We include a sensitivity analysis to examine the impact of including *Aedes* observations from the literature from 1999-2008.

### Climate Data

We obtained climate data on 19 bioclimatic variables in raster form from WorldClim version 2.1 (worldclim.org) which provides averages of data from the years 1970 through 2000. We began our analysis with all 19 variables at 2.5 minute resolution. Raster data was restricted to our defined study area. WorldClim variables describe aspects of either temperature or precipitation-both of which influence mosquito ecology-however, there is no consensus on which variables most effect *Ae. aegypti* distribution. To choose variables to model, we analyzed the variable inflation factor of the variables to assess collinearity (De Marco and Nóbrega 2018). We used the R package usdm (Toxopeus 2014) and selected variables for inclusion in the maximum entropy models which had VIF < 5 (Supplement Table 1.). We chose not to include land cover and population data as climatic variables are assumed to be the main drivers at the continental scale (the scope of this study)(Pearson and Dawson 2003, Bogawski et al. 2019). Additionally, urbanization in West Africa has progressed rapidly during the study period of 2008 to 2020 and may be difficult to capture in a single time-point.

**Table 1.**
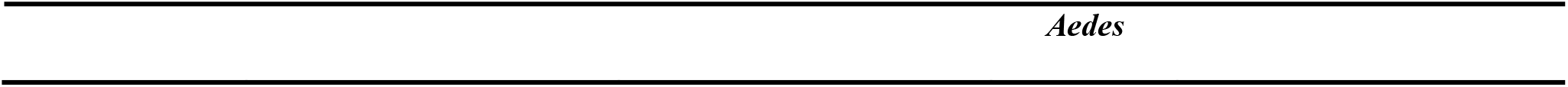
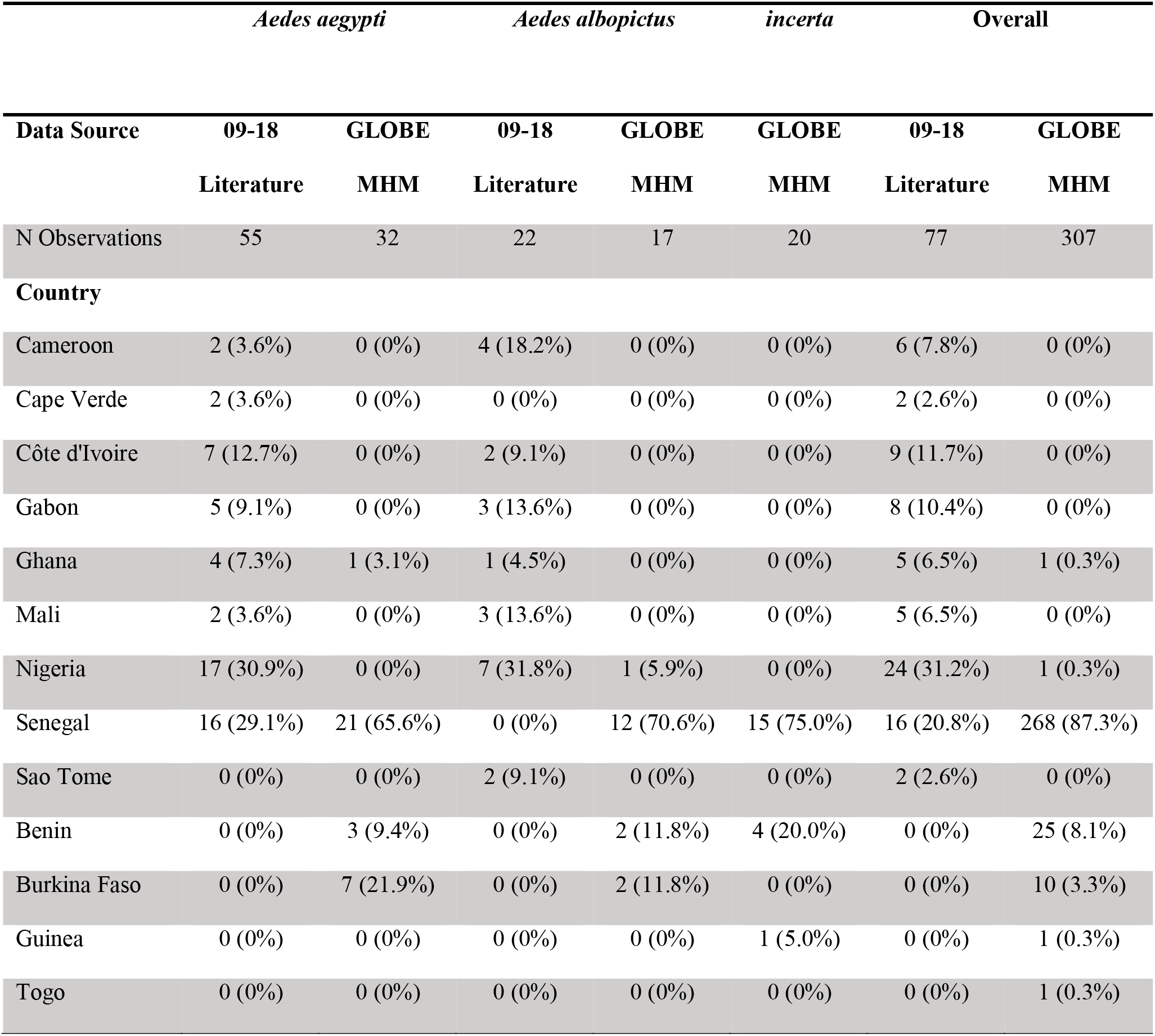
Distribution of *Aedes* mosquito observations in West Africa from literature review and GLOBE Mosquito Habitat Mapper App by species, country, and decade of publication. GLOBE MHM observations were collected 2017 - 2020.

|  | *Aedes aegypti* |  | *Aedes albopictus* |  | *incerta* | Overall |  |
| --- | --- | --- | --- | --- | --- | --- | --- |
| Data Source | 09-18 | GLOBE | 09-18 | GLOBE | GLOBE | 09-18 | GLOBE |
|  | Literature | MHM | Literature | MHM | MHM | Literature | MHM |
| N Observations | 55 | 32 | 22 | 17 | 20 | 77 | 307 |
| **Country** |  |  |  |  |  |  |  |
| Cameroon | 2 (3.6%) | 0 (0%) | 4 (18.2%) | 0 (0%) | 0 (0%) | 6 (7.8%) | 0 (0%) |
| Cape Verde | 2 (3.6%) | 0 (0%) | 0 (0%) | 0 (0%) | 0 (0%) | 2 (2.6%) | 0 (0%) |
| Côte d'Ivoire | 7 (12.7%) | 0 (0%) | 2 (9.1%) | 0 (0%) | 0 (0%) | 9 (11.7%) | 0 (0%) |
| Gabon | 5 (9.1%) | 0 (0%) | 3 (13.6%) | 0 (0%) | 0 (0%) | 8 (10.4%) | 0 (0%) |
| Ghana | 4 (7.3%) | 1 (3.1%) | 1 (4.5%) | 0 (0%) | 0 (0%) | 5 (6.5%) | 1 (0.3%) |
| Mali | 2 (3.6%) | 0 (0%) | 3 (13.6%) | 0 (0%) | 0 (0%) | 5 (6.5%) | 0 (0%) |
| Nigeria | 17 (30.9%) | 0 (0%) | 7 (31.8%) | 1 (5.9%) | 0 (0%) | 24 (31.2%) | 1 (0.3%) |
| Senegal | 16 (29.1%) | 21 (65.6%) | 0 (0%) | 12 (70.6%) | 15 (75.0%) | 16 (20.8%) | 268 (87.3%) |
| Sao Tome | 0 (0%) | 0 (0%) | 2 (9.1%) | 0 (0%) | 0 (0%) | 2 (2.6%) | 0 (0%) |
| Benin | 0 (0%) | 3 (9.4%) | 0 (0%) | 2 (11.8%) | 4 (20.0%) | 0 (0%) | 25 (8.1%) |
| Burkina Faso | 0 (0%) | 7 (21.9%) | 0 (0%) | 2 (11.8%) | 0 (0%) | 0 (0%) | 10 (3.3%) |
| Guinea | 0 (0%) | 0 (0%) | 0 (0%) | 0 (0%) | 1 (5.0%) | 0 (0%) | 1 (0.3%) |
| Togo | 0 (0%) | 0 (0%) | 0 (0%) | 0 (0%) | 0 (0%) | 0 (0%) | 1 (0.3%) |

### Maximum Entropy Modeling

We modeled *Ae. aegypti* distribution using Maximum Entropy (Maxent) version 3.4.4, an approach that creates habitat suitability models by fitting species presence data to climatic variables in raster grid cells (Phillips and DudÍk 2008). The software utilizes the climatic pattern in presence areas to predict other locations with suitable habitat for the study species (Phillips and DudÍk 2008). To adjust for bias in observation data, a bias file was created using the closely related species method (Phillips et al. 2009). All *Aedes* observations including *Ae. aegypti, Ae. albopictus, Ae. incerta*, and unidentified *Aedes* from GLOBE and *Ae. aegypti* and *Ae. albopictus* observations from the literature were projected onto a raster and analyzed using two-dimensional kernel density estimation from the R package MASS to use as the bias layer (Ripley 2002, Hijmans 2020).

Three models were created: 1) using GLOBE MHM data, 2) using literature data from 2009 to 2018, and 3) using GLOBE MHM data combined with literature data from 2009 to 2018. The first two models were used to compare estimated species distributions between models using citizen science data and models using recent data from the literature. The third model was used to determine the change in estimated distribution when adding citizen science data to the literature-derived data. Three additional models were created as a sensitivity analysis, 4) using literature data from 1999 to 2008, 5) using all literature data from 1999 to 2018, and 6) using GLOBE MHM data combined with all literature data (1999 to 2018). We used *Ae. aegypti* observations from 1999-2008 literature as our sensitivity analysis because the time frame in that dataset overlaps with the WorldClim bioclimate variables whereas the data in our main three models do not. The climatic data used was the same for every model. The regularization multiplier (RM) parameter was set to 1.5 to combat potential overfitting issues due to our small datasets. Each model was replicated 20 times and the mean of the replications was examined. All other parameters within Maxent were kept at their default values. Cross-validation compared the runs of the models. Cross-validation used all of the data for validation of models by creating smaller subsets of data as test data for the model that changes each run and using the rest as training data. Models were evaluated based on the area under the receiver operating curve (AUC) for the test data and the prevalence. The AUC is used to estimate how well the model fits the presence and climate data across our study area. The prevalence is the average estimated probability of species present in that area over background sites across all twenty models. Jackknife analysis was run to assess the variable importance of climatic predictor variables for each of the models.

## Results

### Mosquito Presence Data

There were 4,900 observations downloaded from GLOBE Observer’s Advanced Data Access Tool. Of those observations, 3,607 had either larvae, pupae, eggs, or adult mosquitoes present and 307 of those were identified as the genus *Aedes.* Within the *Aedes* identifications, 69 had an accompanying species. After filtering for unique location points, the data contained 13 unique sites for *Ae. albopictus*, 12 unique sites for *Ae. incerta*,and 24 unique sites for *Ae. aegypti. Ae. aegypti* was the only species with sufficient observations to create Maxent models. *Ae. aegypti* were recorded in four countries with most observations (87.3%) located in Senegal (Figure 1).

**Figure 1.**
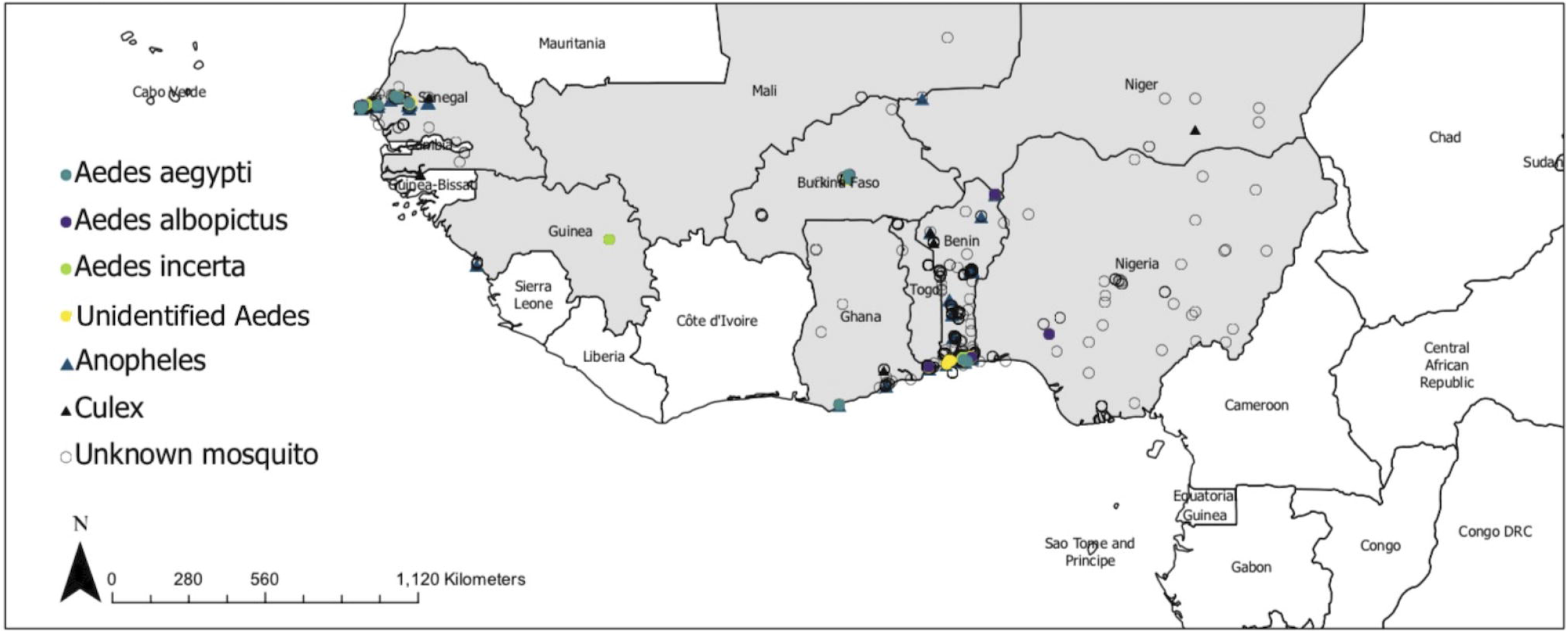
All observations from GLOBE MHM downloaded from the GLOBE Observer Advanced Data Access Tool on November 22nd, 2020.

Senegal was a major contributor of data in the literature as well. Data from literature published from 2009-2018 contained *Aedes* observations from Cameroon, Cape Verde, CÔte d’Ivoire, Gabon, Ghana, Mali, Nigeria, Senegal, and Sao Tome (Figure 2, Table 1). Nigeria and Senegal contained the most Ae. *aegypti* observations with 17 and 16, respectively (Table 1). Literature data published from 1999-2008 contained *Aedes* observations from Burkina Faso, Cameroon, CÔte d’Ivoire, Gabon, Gambia, Ghana, Guinea, Nigeria, Senegal, and Mali (Figure 2).

**Figure 2.**
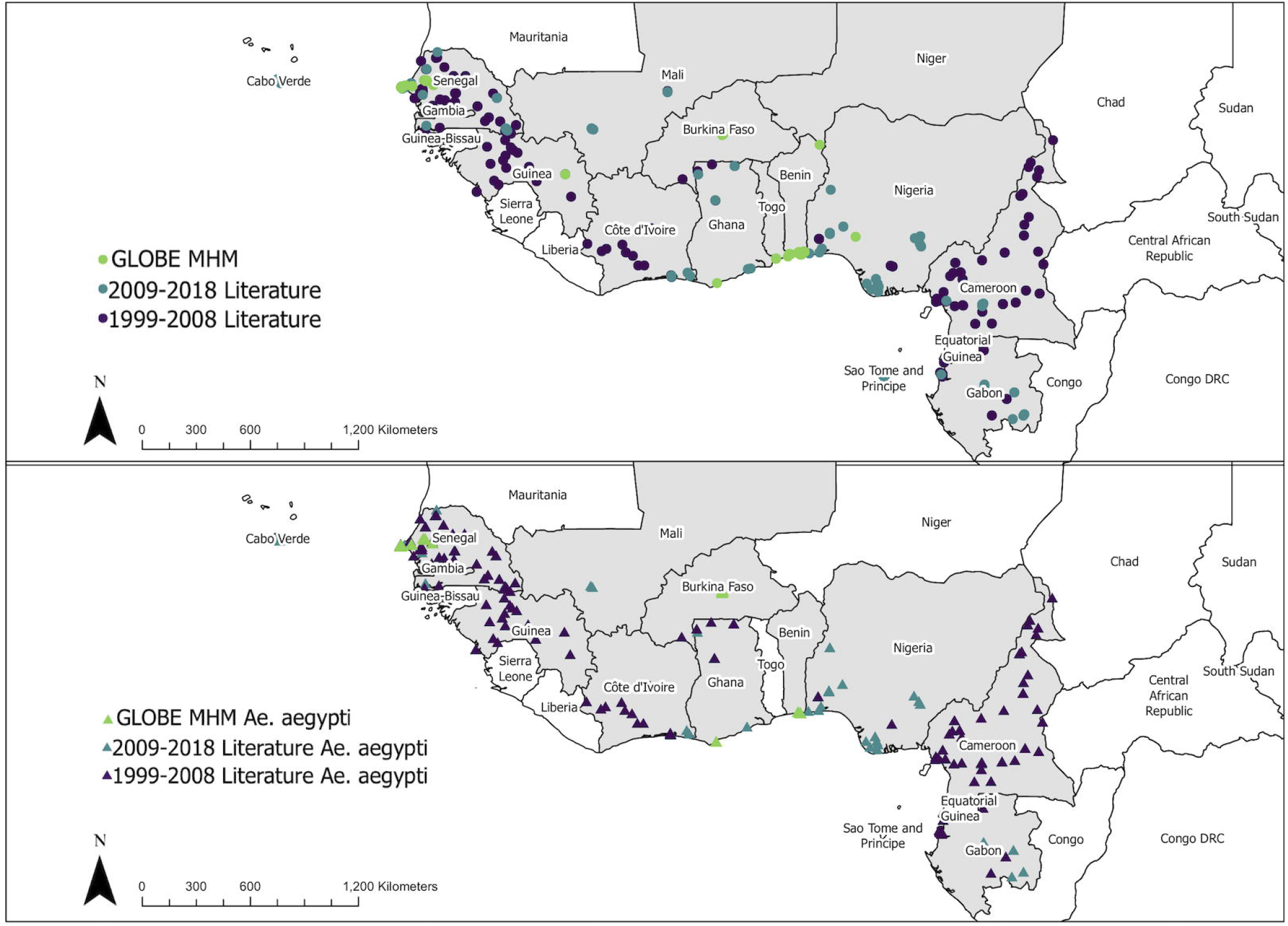
*Aedes* mosquito observations from literature and GLOBE MHM. Top panel: All *Aedes* observations by source dataset. Bottom panel: *Ae. aegypti* observations by source dataset.

### Climate Data

Of the 19 WorldClim bioclimate variables, 8 variables were included in our Maxent models (Figure 3, Supplemental Table 1). Jackknife analysis of climatic predictor variables shows the change in AUC with keeping or removing predictive climatic variables in the models, giving us information about the relative importance that each climatic variable plays in each of the models. The relative importance for each climatic variable varied across the models. In Model (Figure 3A), mean temperature of the wettest quarter (Bio8) lead to the model with the highest AUC when used alone as a predictive variable, whereas precipitation of the coldest quarter (Bio19) lead to the largest decrease in AUC when removed from the full model. In Model 2 (Figure 3B), mean temperature of the coldest quarter (Bioll) alone gave the model with the highest AUC and when removed lead to the largest decrease in predictive accuracy. In Model 3 (Figure 3C), Bio8 led to the highest AUC when used alone and mean diurnal range (Bio2) led to the greatest decrease in AUC when removed.

**Figure 3.**
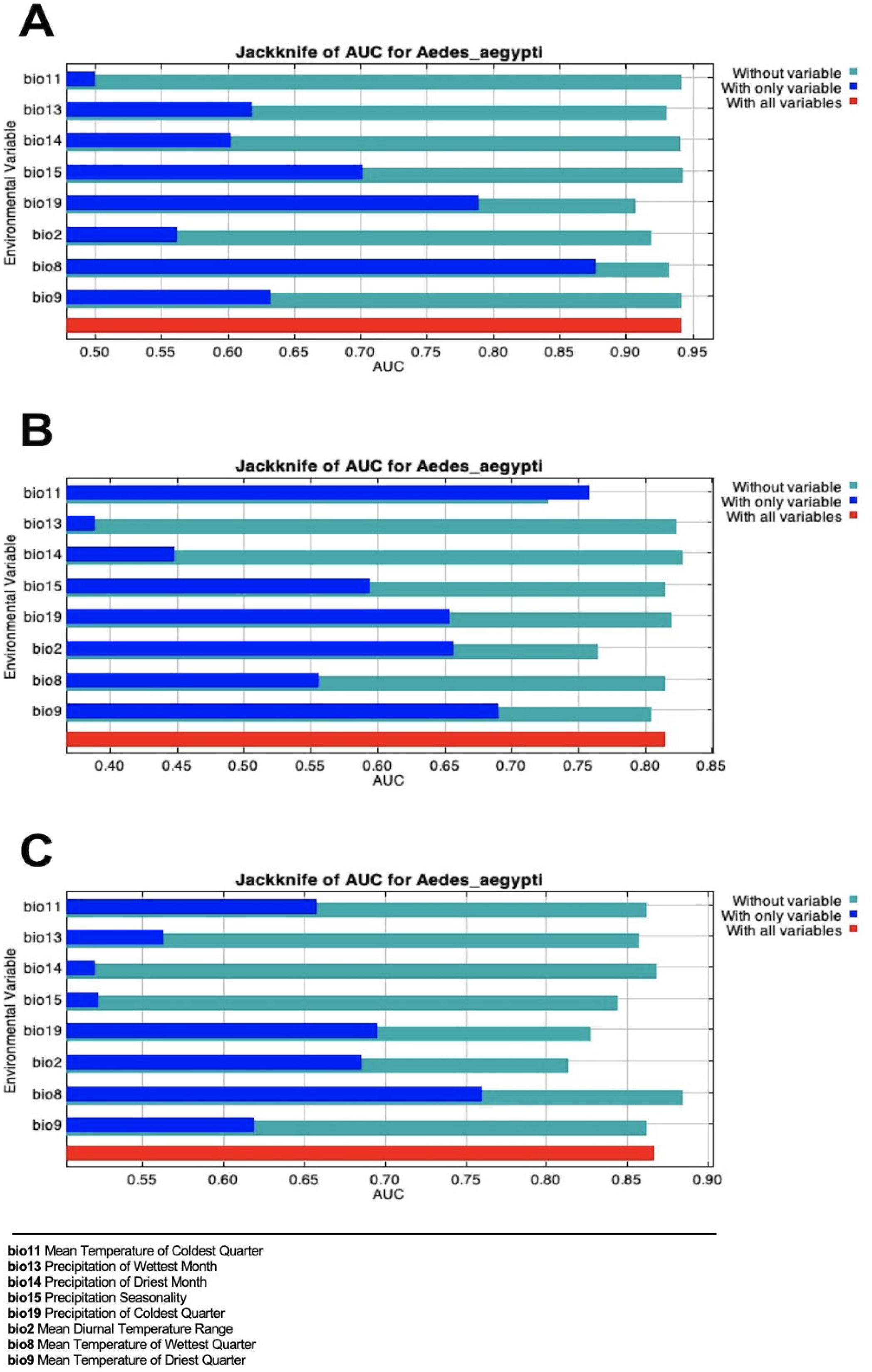
A: Jackknife plot of climatic variables for Model 1; Maximum Entropy output using GLOBE MHM data. B: Jackknife plot of climatic variables for Model 2; Maximum Entropy output using 2009-2018 literature data. C: Jackknife plot of climatic variables for Model 3; Maximum Entropy output using a combination of GLOBE MHM data and 2009-2018 literature data.

### Maximum Entropy Modeling

The suitable habitat predicted by each model varied in prevalence (Table 2). Model 1, using data from GLOBE MHM estimated average prevalence relatively lower than models that used literature data, with predicted high prevalence of *Aedes aegypti* in Senegal and scattered presence across southern Mali, Burkina Faso, and Nigeria (Figure 4A). Model 2, using the 2009-2018 literature data (Figure 4B) predicted widespread suitable habitat for *Ae. aegypti*, with high presence probability estimated for-an area including the coast of CÔte d’Ivoire, Ghana, Togo, Benin, Nigeria, Cameroon, and Gabon. Areas across Senegal and Burkina Faso were also predicted to have high probability of *Ae. aegypti* presence. Model 3 (Figure 4C) used data combined from the GLOBE MHM and the 2009-2018 literature data and predicted a lower average prevalence than Model 2, but the test AUC increased compared to Model 2. Using the combined dataset, Maxent predicted more Ae. *aegypti* habitat suitability than Model 1, covering regions in Senegal, Burkina Faso, Nigeria, and along the coasts of CÔte d’Ivoire, Ghana and Togo.

**Table 2.** Model output for prevalence of *Ae. aegypti*, area under the curve (AUC) of the ROC, and the standard deviation of the AUC for all combinations of data ran in Maxent. Prevalence describes the proportion of the area modeled estimated to be suitable for *Ae. aegypti* and *Ae. spp.* AUC and AUC standard deviation are reported for the test data in each model.

| Model | Data Source | N Observations | Prevalence | Test AUC | Test AUC SD |
| --- | --- | --- | --- | --- | --- |
| 1 | GLOBE MHM | 24 | 0.1901 | 0.9417 | 0.056 |
| 2 | 09-18 Literature | 55 | 0.3713 | 0.8157 | 0.167 |
| 3 | GLOBE MHM and<br>09-18 Literature | 79 | 0.293 | 0.8669 | 0.109 |
| 4 | 99-08 Literature | 179 | 0.1693 | 0.7631 | 0.078 |
| 5 | 99-18 Literature | 234 | 0.2374 | 0.7124 | 0.0838 |
| 6 | GLOBE MHM and<br>99-18 Literature | 258 | 0.1999 | 0.7601 | 0.068 |

**Figure 4.**
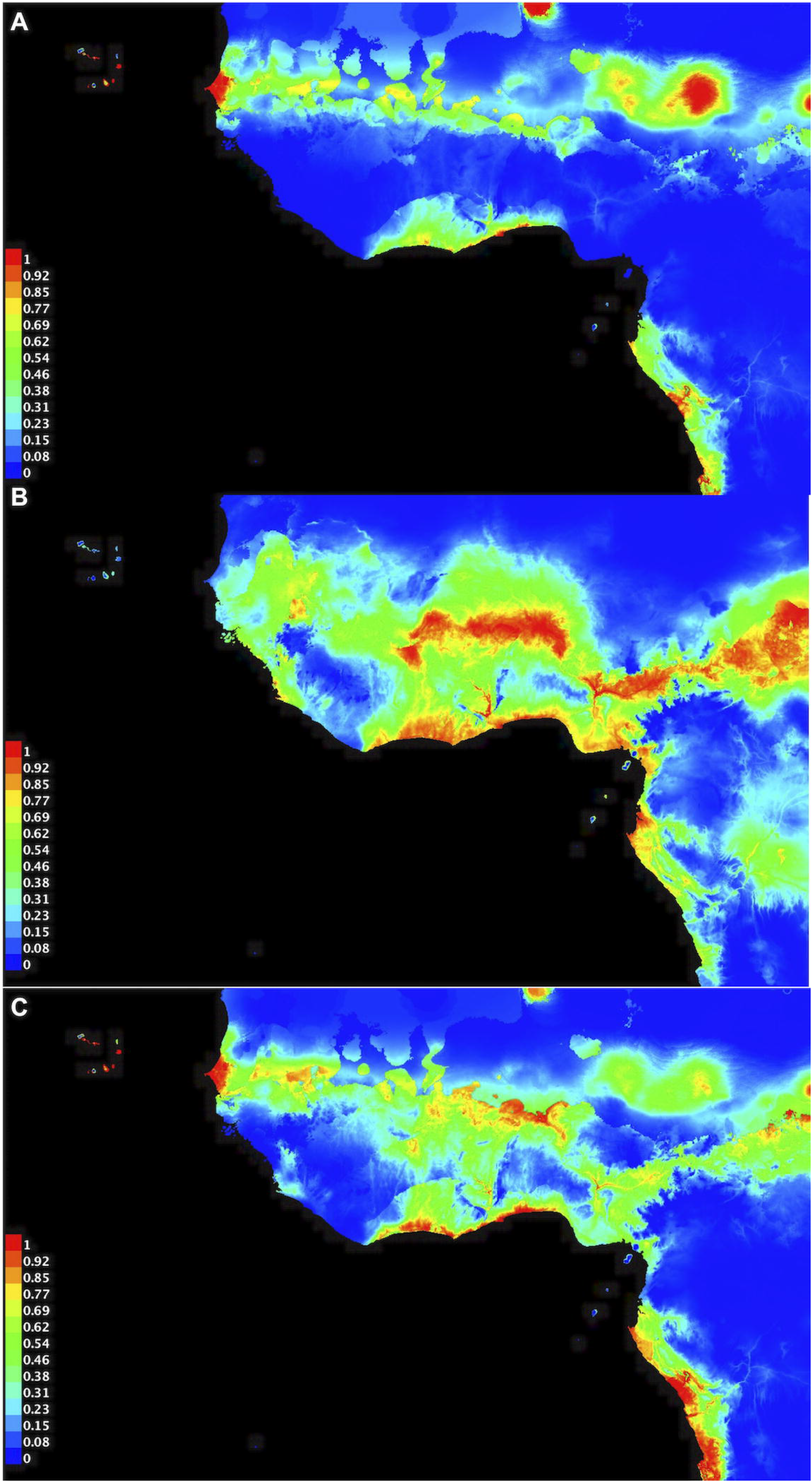
Estimated *Ae. Aegypti* habitat suitability in West Africa using Maximum Entropy Modeling. A) Models fit to *Ae. Aegypti* observations from GLOBE MHM; B) Models fit to data from the literature published from 2009 to 2018; C) Models fit to data from GLOBE MHM and the 2009-2018 literature.

### Sensitivity Analyses

The data from literature published 1999-2008 was the largest individual data set (n=179), however Model 4 both predicted lower prevalence of *Ae. aegypti* and had a lower AUC than any of Models 1-3. Model 5 (Supplement figure 1B), using all literature data published from 1999 to 2018, similarly estimated lower prevalence than from the model including only recent data and had decreased fit to the data. Model 6 (Supplemental Figure 1C), combining all *Ae. aegypti* data from the literature and from GLOBE MHM, similarly estimated decreased prevalence compared to Model 2 and had decreased fit to the data (Table 2).

## Discussion

We compared predictions of *Ae. Aegypti* mosquito distribution in West Africa from models using literature-derived and citizen science-derived data. Addition of citizen science data improved the model fit to the data, suggesting an ongoing role for citizen science in improving our understanding of *Aedes* mosquito distribution in the region. However, all of our models were likely subject to location bias, highlighting the need for increased surveillance of *Aedes* species mosquitoes throughout West Africa. Despite the lack of observations and potential location bias in the data, all models including observations from the literature predicted widespread distribution of *Ae. aegypti* in the region.

Previous global mapping efforts have estimated the distribution of *Ae. aegypti* in West Africa using a variety of methods(Kraemer et al. 2015, Gloria-Soria et al. 2016, Kamal et al. 2019). All our models estimated lower prevalence of *Ae. aegypti* in West Africa compared to previous global models of *Ae. aegypti.* Few presence points in our study may have contributed to the lower prevalence values that we found (van Proosdij et al. 2016). However, all previous models have been at a wider spatial scale (Kraemer et al. 2015, Gloria-Soria et al. 2016, Kamal et al. 2019, Ryan et al. 2019). To our knowledge, this study is the first model of *Ae. aegypti* distribution focusing solely on the region of West Africa. Aedes behavior and habitat selection may vary regionally, due to subspecies variation, local-level competition, or other factors-potentially biasing estimates from global models.

Citizen science can be used to increase *Aedes* surveillance data, but like any surveillance method, comes with limitations including recruitment, adherence, and concerns of data quality. Recruitment of citizen scientists to participate in surveillance projects is a challenge (Basso et al. 2015, Lwin et al. 2017), with participation unequal across socio-economic groups (Hobbs and White 2012). Lack of adherence to protocols may occur among citizen scientists (Cohnstaedt et al. 2016, Johnson et al. 2018), however similar problems can occur in traditional mosquito surveillance. There are known data biases in GLOBE MHM; participants only identified mosquito genus 14% of the time and 7.8% of observations from 2016-2019 were flagged as problematic previously (Amos et al. 2020). Some species may have been misclassified within the citizen science data used. Some data put into GLOBE MHM has been verified by scientists at organizations partnering with GLOBE, but much of the identification of mosquito genus and species in the app relies on morphological identifications by untrained citizen scientists. GLOBE MHM attempts to mitigate this bias by walking the user through a series of larval identification. The technique presented requires the user to have a magnifying lens attachment for their smartphone camera, which not all users will have or use properly. Misidentification of species is not accounted for within Maxent and this may have influenced model estimates (Lozier et al. 2009).

Although there was likely bias created from GLOBE MHM observations in general and particularly due to the overweighting of observations from Senegal, the long-standing relationships between schools and the GLOBE program is thought to improve the quality of the data. Of the observations included in this study, 81% (N = 249) were recorded by lycee Thumakha, GLOBE’s largest school partner in Senegal. They partnered with GLOBE in 2015 and have 100 students participating in the <u>program</u>. Future research should focus on investigating the accuracy of GLOBE Observer data in Senegal alone where data may be sufficient to examine mosquito distribution on a local-level scale. Building similar partnerships throughout West Africa will improve data from citizen science projects, as well as expand local education and knowledge of the risks and prevention methods to control vector-borne diseases.

Multiple methods have been utilized for engaging citizen scientists in mosquito surveillance. Some have focused on collection and submittal of a biological specimen (Cohnstaedt et al. 2016, Walther and Kampen 2017), which are then identified by trained entomologists. Other projects have utilized citizen scientists to quantify mosquitoes caught in surveillance equipment to aid researchers in gathering data (Bazin and Williams 2018, Johnson et al. 2018). Recently, mobile applications have emerged as a popular tool for citizen science-based mosquito surveillance (Lwin et al. 2017, Palmer et al. 2017, de Souza Silva et al. 2018) as applications can increase the accessibility of mosquito surveillance due to widespread smart phone use. Such applications have demonstrated mixed success as a surveillance tool (Lwin et al. 2017, Eritja et al. 2019). Use of the Mosquito Alert application (Palmer et al. 2017) led to the first detection of *Aedes japonicus* in Spain (Eritja et al. 2019). Others have been less successful due to inherent design issues (Lwin et al. 2014, Lwin et al. 2016, Lwin et al. 2017).

We recently introduced GLOBE to Ghanaian students at Junior High Schools near Accra. While there were some difficulties with implementation, including a lack of compatibility with widely-used varieties of tablets and phones available in low-resource settings, at the end of the study, the teachers and students were enthusiastic about the program and interested in seeing it continues. Following the training and data collection, students understood their role in mosquito control as a way to reduce mosquito-borne diseases, particularly *Aedes*-borne diseases. Teachers valued the hands-on instruction in the app and expressed enthusiasm for integrating education on mosquitoes and arboviruses in the schools. The ability for students to do “field work” in their own back yards was engaging and highlighted the potential for the GLOBE program to expand and contribute to mosquito surveillance in the future.

The estimates of *Ae. aegypti* mosquito distribution in West Africa produced herein are subject to a number of important limitations: Observations used in these models were locationally biased due to clustering in the datasets used. Location bias may have biased estimates of environmentally suitable habitats as well as have led to over-fitting of models with few presence points. While we attempted to adjust for location bias with both bias file offsets and tweaking the RM parameter, this may not have been sufficient. We additionally were unable to adjust for land cover due to fluctuations in land cover over the (long) study period. Future estimates may benefit from using predicted land cover data to account for rapid urbanization. Additionally, while WorldClim is used widely in Maxent models (Lubinda et al. 2019, Tiffin et al. 2019, Ibáñez-Justicia et al. 2020), the dates of the WorldClim data (1970-2000) do not overlap with the dates of the *Aedes* presence data used in Models 1-3. Climate change has been noted on a short time scale-2000 to 2009-in West Africa (Barry et al. 2018) and in such a dynamic region, both mosquito distribution and local climate patterns are likely in constant flux, requiring real-time data to detect changes. Finally, we note that growing evidence suggests *Ae. albopictus* also plays an important role in arboviral infections in the region, however there was not sufficient data to examine *Ae. albopictus.* distribution

## Conclusions

There is a need for expanded *Aedes* mosquito surveillance throughout West Africa. Despite some heterogeneity in implementation and though quality of data collected by citizen scientists is a concern, these issues are outweighed by the potential benefits of increased data in mosquito-borne disease research. The models presented in this manuscript suggest data from these mobile applications can be useful in supplementing our understanding of vector distribution. While both traditional mosquito surveillance and citizen science have pros and cons, using them in combination can improve understanding of mosquito distributions and *Aedes*-borne disease risk.

## Acknowledgements

We would like to thank all schools and citizen scientists who contributed mosquito observations to the GLOBE MHM. This work was funded by a grant through the GLOBE Engaging Citizens in the Forecasting and Observation of Mosquito Threats Program from the United States Department of State and the University Corporation for Atmospheric Research (SLMAQM17CA2074). The GLOBE Implementation Office (GIO) managed the activities in the citizen and mosquito program. The opinions, findings and conclusions stated herein are those of the authors and do not necessarily reflect those of GIO, UCAR, or the US State Department.

## Data Availability Statement

All data and code used in this manuscript are available at https://github.com/eafreeman/aedesglobe.

---

**Supplemental Table 1.** Variable descriptions of all environmental variables in WorldClim dataset and examined in variable inflation factor analysis. Variables were chosen to be used in the Maxent models if they had a variable inflation factor of under 5.

| Variable | Description | Selected for Maximum Entropy Modeling |
| --- | --- | --- |
| Bio1 | Annual Mean Temperature | No |
| Bio2 | Mean Diurnal Range (Mean of monthly (max temp = min temp)) | Yes |
| Bio3 | Isothermality (Bio2/Bio7)x100 | No |
| Bio4 | Temperature Seasonality (Standard Deviation x 100) | No |
| Bio5 | Max Temperature of Warmest Month | No |
| Bio6 | Min Temperature of Coldest Month | No |
| Bio7 | Temperature Annual Range (Bio5-Bio6) | No |
| Bio8 | Mean Temperature of Wettest Quarter | Yes |
| Bio9 | Mean Temperature of Driest Quarter | Yes |
| Bio10 | Mean Temperature of Warmest Quarter | No |
| Bio11 | Mean Temperature of Coldest Quarter | Yes |
| Bio12 | Annual Precipitation | No |
| Bio13 | Precipitation of Wettest Month | Yes |
| Bio14 | Precipitation of Driest Month | Yes |
| Bio15<sup>+</sup> | Precipitation Seasonality (Coefficient of Variation) | Yes |
| Bio16 | Precipitation of Wettest Quarter | No |
| Bio17 | Precipitation of Driest Quarter | No |
| Bio18 | Precipitation of Warmest Quarter | No |
| Bio19 | Precipitation of Coldest Quarter | Yes |

+ The coefficient of variation for precipitation seasonality is the ratio of the standard deviation of the monthly total precipitation to the mean monthly total precipitation expressed as a percentage (Hijmans et al. 2005).

**Supplement Figure 1.**
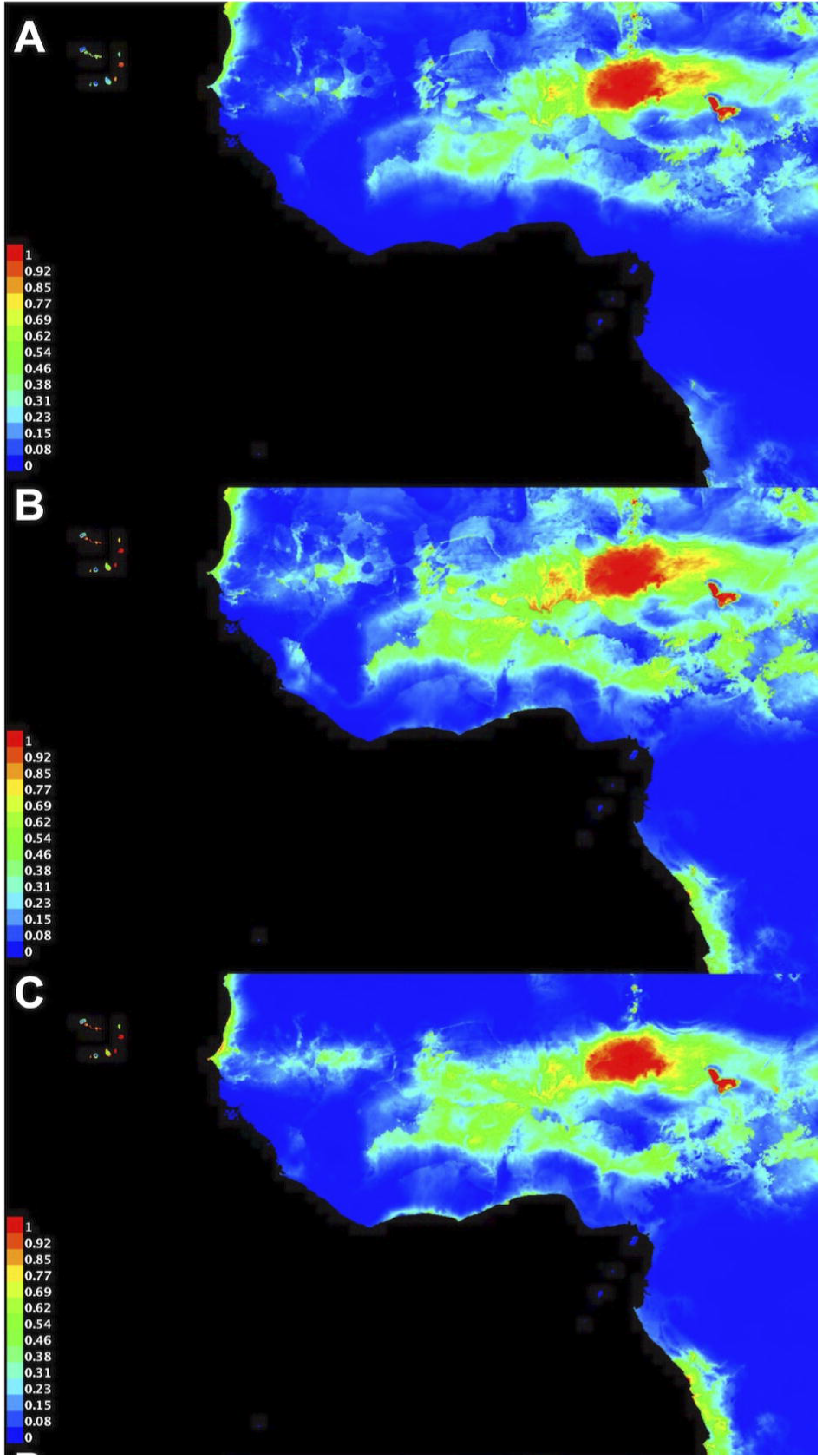
**A.** Model 4; Maximum Entropy output of *Ae. aegypti* habitat suitability in West Africa using data from 1999-2008 literature data. **B.** Model 5; Maximum Entropy output of *Ae. aegypti* habitat suitability in West Africa using 1999-2018 literature data, **C.** Model 6; Maximum Entropy output of *Ae. aegypti* habitat suitability in West Africa using a combination of GLOBE Observer app data and 1999-2018 literature data.

## Notes

https://github.com/eafreeman/aedes_globe

### Competing Interest Statement

The authors have declared no competing interest.

